# Life-history traits, pace of life and dispersal among and within five species of *Trichogramma* wasps: a comparative analysis

**DOI:** 10.1101/2023.01.24.525360

**Authors:** Chloé Guicharnaud, Géraldine Groussier, Erwan Beranger, Laurent Lamy, Elodie Vercken, Maxime Dahirel

## Abstract

Major traits defining the life history of organisms are often not independent from each other, with most of their variation aligning along key axes such as the pace-of-life axis. We can define a pace-of-life axis structuring reproduction and development time as a continuum from less-fecund, longer-developing ‘slow’ types to more-fecund, shorter-developing ‘fast’ types. Such axes, along with their potential associations or syndromes with other traits such as dispersal, are however not universal; in particular, support for their presence may be taxon and taxonomic scale-dependent. Knowing about such life-history strategies may be especially important for understanding eco-evolutionary dynamics, as these trait syndromes may constrain trait variation or be correlated with other traits. To understand how life-history traits and effective dispersal covary, we measured these traits in controlled conditions for 28 lines from five species of *Trichogramma*, which are small endoparasitoid wasps frequently used as a biological model in experimental evolution but also in biocontrol against Lepidoptera pests. We found partial evidence of a pace-of-life axis at the interspecific level: species with higher fecundity also had faster development time. However, faster-developing species also were more likely to delay egg-laying, a trait that is usually interpreted as “slow”. There was no support for similar covariation patterns at the within-species line level. There was limited variation in effective dispersal between species and lines, and accordingly, we did not detect any correlation between effective dispersal probability and life-history traits. We discuss how expanding our experimental design by accounting for the density-dependence of both the pace of life and dispersal might improve our understanding of those traits and how they interact with each other. Overall, our results highlight the importance of exploring covariation at the “right” taxonomic scale, or multiple taxonomic scales, to understand the (co)evolution of life-history traits. They also suggest that optimizing both reproductive and development traits to maximize the efficiency of biocontrol may be difficult in programs using only one species.

## Introduction

Life history describes the life cycle of an organism, how fast and how much it grows, reproduces, and survives. It is the direct product of a collection of phenotypic traits, called life-history traits (Flatt & Heyland, 2011). Those traits include growth and mortality rates, survival, reproductive investment or even the lifespan, and can be age- or stage-specific. When all life-history traits and the values they can take are combined, many pathways can lead to evolutionary success, resulting in the high diversity of what are called life-history strategies, the covariation through time and space of different traits, found across the tree of life. This high diversity can be observed at multiple taxonomic levels, from the phylum level to within species (Gaillard et al., 1989; Healy et al., 2019; Olsen et al., 2018). Yet, resource limitation means not all strategies are possible: indeed, analyses of life-history traits across taxa and hierarchical levels often reveal that a large part of the variation in organisms’ life histories can be summarised on a small number of key axes, which often reflect trade-offs between life-history components. It is generally accepted that those life-history-trait correlations arise from trade-offs between allocating a certain amount of acquired resources into one trait or another, with limitations arising from a limited pool of resource to draw from, physiological constraints, and from the influence of the environment, resulting in a variety of strategies maximizing fitness (Laskowski et al., 2021; Stearns, 2000).

One specific axis has been termed the pace of life and corresponds to a correlation between life-history traits sorting organisms along a fast-slow continuum (Braendle et al., 2011; Stearns, 1983). Many trait combinations can be used to characterize a pace-of-life axis (Gaillard et al., 2016), discriminating low reproduction, long development and long lifespan (slow types) on one side from high reproduction, short development, and short lifespan (fast types) on the other. Pace-of-life axes have been identified in multiple comparative analyses across taxonomic ranks (Auer et al., 2018; Healy et al., 2019; Williams et al., 2010) although the traits that cluster to form this axis are not always the same (Bielby et al., 2007). But despite its conceptual appeal and simplicity, the pace-of-life axis should not be assumed as the one unique axis structuring life histories: the proportion of variance explained by such an axis varies between taxa (Healy et al., 2019), and in many cases, alternative axes structuring variation also emerge (Bakewell et al., 2020; Mayhew, 2016; Wright et al., 2020). Moreover, it seems the narrower the taxonomic focus (from tree of life-wide analyses to within-species comparisons), the harder it is to find the presence of a pace of life, and the way life-history variation is structured in one species/taxon cannot always be generalized to others. Adding complexity to the correlations of life-history traits, the pace-of-life syndrome hypothesis supposes that the pace of life can co-evolve with one or many other phenotypic traits. They can be physiological (Auer et al., 2018; Ricklefs & Wikelski, 2002), behavioural (Réale et al., 2010; Wolf et al., 2007), or associated with other traits like dispersal.

Dispersal can be described as any movement potentially leading to a flux of genes or individuals across space (Ronce, 2007), and is a key component influencing both ecological and evolutionary dynamics, so much that it is sometimes described as a life-history trait in its own right (Saastamoinen et al., 2018). Dispersal often covaries with other traits, including other life-history traits (Clobert et al., 2012), in so-called dispersal syndromes (Ronce, 2012). Dispersal syndromes have been observed and compared at multiple taxonomic levels, both across (Stevens et al., 2012, 2014) and within species (Jacob et al., 2019). Therefore, it is not surprising that many works have been dedicated to the integration of dispersal along the main life-history axes, and the derivation of ecological and evolutionary implications. This includes, for instance, the idea of a trade-off between competition and colonization where species that are good at colonizing, with high fecundity or dispersal, are in return poor competitors between or among species (e.g. Calcagno et al., 2006; Yu & Wilson, 2001), and other studied links between dispersal and fecundity (Bonte & De La Peña, 2009; Crossin et al., 2004; Gu et al., 2006; Karlsson & Johansson, 2008). Rather than idiosyncratic correlations between dispersal and specific life-history traits, the pace-of-life syndrome hypothesis suggests dispersal, among others, to be a risky trait linked to the pace of life itself (Cote, Clobert, et al., 2010; Réale et al., 2010). In plants for instance, there is a relation between seed dispersal abilities and the fast-slow continuum, where a high capacity to disperse is correlated with faster life histories at the species level (Beckman et al., 2018). While many studies found a positive correlation between the pace of life and short-scale movement, like the exploration of a continuous patch, or the activity level within an arena (Gangloff et al., 2017; Lartigue et al., 2022; Rádai et al., 2017), directly transposing short-scale conclusions like exploration or activity, to longer-scale metrics, like dispersal rates or the decision to disperse in discrete landscapes, is not always relevant (Cote, Fogarty, et al., 2010; Harrison et al., 2015; Pennekamp et al., 2019). Dispersal/life-history syndromes can lead to different ecological and evolutionary results from when traits are considered as independent. Correlation between traits, but also the strength or shape of this relationship can impact both the ecological and evolutionary dynamics of a population (Maharjan et al., 2013; Ochocki et al., 2020).

In that context, we explored first the presence of a pace-of-life axis and then the relationship between the pace of life and effective dispersal in five species of *Trichogramma* wasps. *Trichogramma* (Hymenoptera: Trichogrammatidae) are small (< 1 mm when adult) parasitoids that develop inside the eggs of their hosts, mainly Lepidoptera. They are also model species in ecology and invasion biology studies thanks to their small size, rather short development time (13-15 days, at 22 °C), or also the fact that lines can be either sexual or asexual. The goal of this study is therefore to improve our knowledge of life-history trade-offs specifically in *Trichogramma* for future studies of eco-evolutionary dynamics, but also more generally in insects, which are under-represented in both pace-of-life (but see Blackburn 1991) and pace-of-life syndromes studies (38 invertebrate species vs 141 vertebrates in Royauté et al., 2018). Potential reasons for this under-representation include a lack of data (Bakewell et al., 2020) or the difficulty to study and compare insect parasitoids, as their life-history traits are also subject to their host ecology (Mayhew, 2016). Using lab-reared lines belonging to five species of *Trichogramma*, we measured female fecundity, effective dispersal, and development time under experimental conditions, and analysed the line- and species-level covariation between these traits using multivariate Generalized Linear Mixed Models (Careau & Wilson, 2017; Dingemanse & Dochtermann, 2013). While this study is mostly exploratory, we can make some hypotheses: based on previous experiments that analysed trait variation between *Trichogramma* lines (Lartigue et al., 2022), or species (Özder & Kara, 2010), we can expect to observe trade-offs between fecundity and development time at the interline or interspecies level. In addition, as a relationship was found between activity and fecundity in Lartigue et al. (2022), there is a possibility that one or several life-history traits are linked to effective dispersal in a dispersal syndrome at a species or line level.

## Materials and methods

### Biological material

*Trichogramma* are endoparasitoids, which means that females lay their eggs inside their hosts, where the larvae will develop by feeding on the host and ultimately killing it, as opposed to ectoparasitoids, who lay their eggs and develop outside their host. As some of *Trichogramma* hosts are Lepidopteran pest species, several *Trichogramma* species are used as biological control agents, and have shown to work well (Smith, 1996). For instance, *T. brassicae* is used on a large scale against *Ostrinia nubilalis*, the European corn borer (Mertz et al., 1995), and *T. evanescens*, *T. cacoeciae,* or a mix of the two species can be used against *Cydia pomonella*, an apple pest (Sigsgaard et al., 2017). In addition to their interest as laboratory model species to investigate the pace of life, the identification of correlations between life-history traits in *Trichogramma* could open up new avenues to improve their efficiency as biocontrol agents, through the optimization of their rearing or field performance (Akbari et al., 2012; Consoli et al., 2010).

For this experiment, 32 different lines of *Trichogramma* were originally selected among the collection from the Biological Resource Center (BRC) “Egg Parasitoid Collection”(CRB EP-Coll, Sophia Antipolis; Marchand et al., 2017). We restricted our choice to the only five sexual species where at least three lines were available. Within each species, we selected at random at least three lines per species and up to ten, with a total target of 32 lines for feasibility. Four lines did not correctly synchronize during preparation and could not be used, resulting in 28 lines in the actual experiment (**Table 1**). The Biological Resource Center rears lines on eggs of the Mediterranean flour moth *Ephestia kuehniella* (Lepidoptera: Pyralidae) at 18 °C, 70 % ± 10 % relative humidity, L:D 16:8. Most lines were founded from a single original clutch each, mostly collected between 2013 and 2016 in different parts of France, and one line comes from a crossing of three single-clutch lines made in 2019 (**Supplemental Table S1-1**). With approximately 15 generations per year under those rearing conditions, lines from the BRC collection are expected to have a very low genetic variance at the time of the experiment (as seen for *Trichogramma brassicae* in the supplemental material of Dahirel, Bertin, Haond, et al., 2021). Little is known about the genetic diversity in the wild, but it is expected to be low as a survey in France and Spain collected only two to three haplotypes for *T. evanescens*, *T. semblidis* and *T. brassicae* (Muru, 2021). After collecting the lines from the BRC, we kept them on *E. kuehniella* eggs at 22 °C, 70 % ± 10 % relative humidity, L:D 16:8 for two to three generations before starting the experiment. Host eggs were irradiated with UV for 15 minutes before use; this sterilization method kills the embryo while keeping it viable as a wasp host (St-Onge et al., 2014). Each female used for the experiment was isolated randomly from the rest of the population 24 hours after emerging, as *Trichogramma* start mating as soon as individuals emerge from host eggs (Doyon & Boivin, 2006). Therefore, all females during the experiment were between 24 to 48 hours old.

**Table 1:** Summary of the *Trichogramma* species and lines used in the experiment, among the total number of lines available in the BRC at the time.

| Species | Species authority | Number of lines used (number available in the BRC) |
| --- | --- | --- |
| *T. bourarachae* | Pintureau & Babault, 1988 | 4 (4) |
| *T. brassicae* | Bezdenko, 1968 | 9 (22) |
| *T. evanescens* | Westwood, 1833 | 7 (21) |
| *T. principium* | Sugonjaev & Sorokina, 1976 | 3 (4) |
| *T. semblidis* | (Aurivillius 1898) | 5 (5) |

### Experimental design

We used both single- and two-vial systems to measure life-history traits (**Fig. 1**). In single-vial systems (12 replicates per line), we placed one randomly selected mated *Trichogramma* female between 24 to 48 hours old into a plastic vial (5 cm diameter, 10 cm height). We also added a non-limiting quantity of irradiated *Ephestia kuehniella* eggs on a paper strip (hundreds of host eggs in approximatively 1.4 × 1 cm, see **Supplemental Figure S2-1**). This system was used to measure development time and fecundity traits. In two-vial systems (20 replicates per line), the setup was similar to the previous one, with the exception that a see-through 40 cm long plastic pipe (5 mm of internal diameter, large enough for species of less than a millimetre in size) connected the first vial (where the wasp was deposited) to another one with the same dimensions, also containing a non-limiting quantity of irradiated eggs. The ends passed through the centre of the foam plugs without protruding from them. While little is yet known about how females locate host eggs (Consoli et al., 2010), this setup was inspired by previous studies on experimental expansions on *Trichogramma* (Dahirel, Bertin, Calcagno, et al., 2021; Dahirel, Bertin, Haond, et al., 2021) and allowed us to estimate effective dispersal probability in conditions similar to previous experimental expansions. Even though fecundity and development-time data could also be collected in this second setup, we refrained from analysing them here due to the complexities of accounting for the effects of dispersal and dispersal costs, compared to the single-vial setup. Females were left in those vials for 48 h under standardized conditions: 22 °C, 70 % relative humidity, L:D 16:8. After 48 h, the egg strips were isolated in glass vials (1 cm diameter, 4 cm height), and kept under the same standardized conditions. Please note that even if plasticity can be observed in *Trichogramma* (Krishnaraj, 2000; Pinto et al., 1989), we focused our study on the presence or not of a pace-of-life under the standard conditions used in experimental expansions on *Trichogramma*, allowing us to make more direct links between our results in this study and future results in experimental expansions.

**Figure 1:**
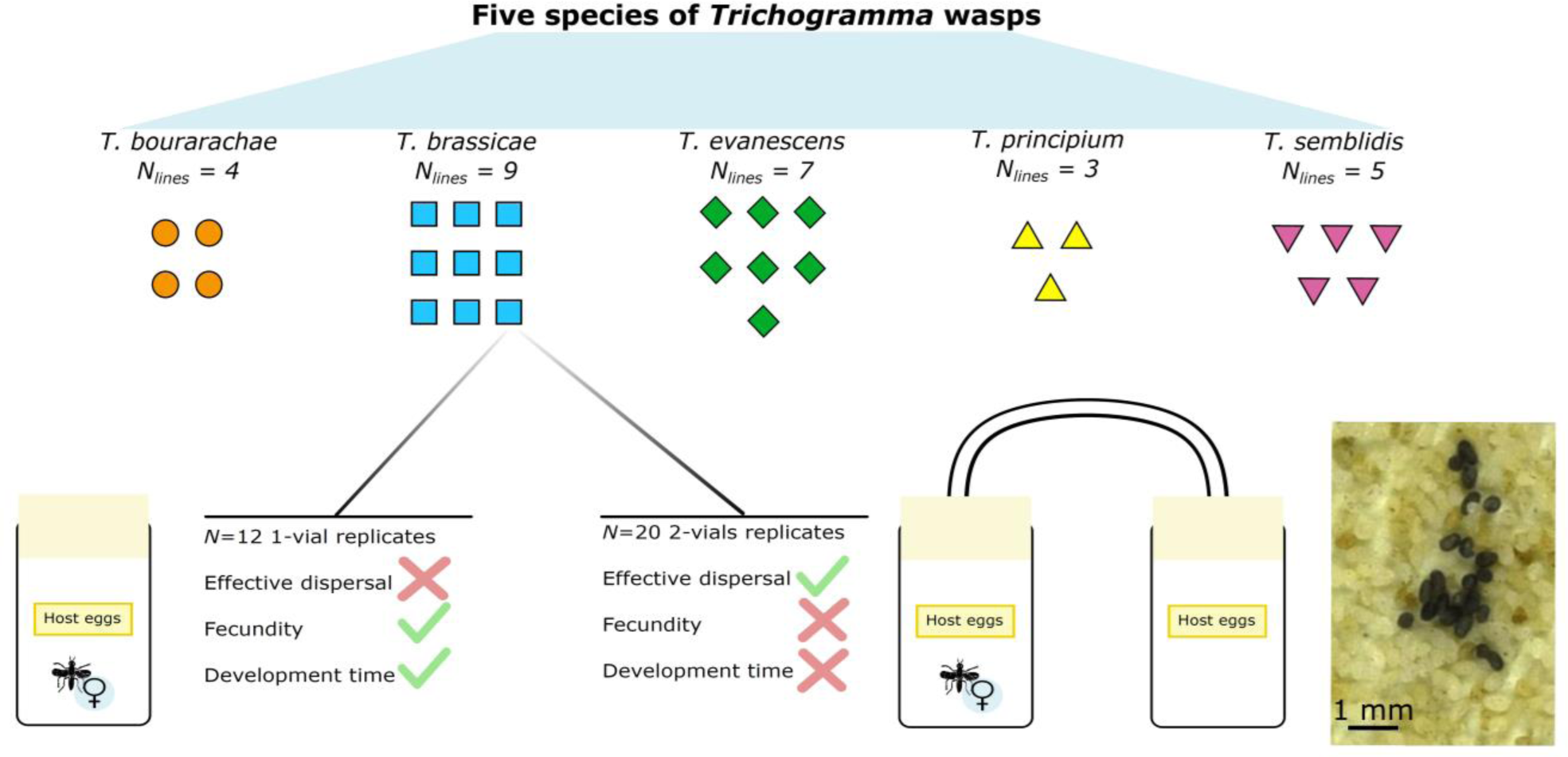
Summary of the experimental design used for measuring fecundity, effective dispersal, and development time. Inset (bottom right): picture of; parasitized host eggs, in black, easily visible among the off-white unparasitized hosts, one week after the experiment.

### Phenotyping

For endoparasitoids, the body size is highly dependent on the host size. In our case, all species were maintained and experimented using *E. kuehniella* as host eggs, which are small enough to allow only one viable descendent (Corrigan et al., 1995) and were provided in high enough quantity to avoid superparasitism (as multiple eggs within one host might affect the viable descendent size). Therefore, we assumed that size variance was probably highly limited, with little to no correlations between hind tibia length (one proxy of individual size) and other traits (Pavlík, 1993) and did not measure size.

### Fecundity and dispersal

A week after isolation, parasitoid larvae were developed enough to blacken the host egg, allowing the visual identification of successfully parasitized eggs (picture in **Figure 1**). Egg strips (one for single vial, two for two-vial systems) were then photographed (resolution: 6016 × 4016 pixels, for a real field of view size of around 12 × 8 cm) using a Nikon D750 camera (lens: AF-S Micro NIKKOR 60 mm f/2.8 G ED) fixed above the strips.

Fecundity was measured by manually counting the number of blackened eggs in each picture using ImageJ (Schneider et al., 2012). Even though superparasitism (more than one parasitoid egg laid per host) is frequent for *Trichogramma*, it tends to be avoided when an unlimited number of unparasitized eggs are present for single females (in *T. chilonis*, Wang et al., 2016). As in Özder & Kara (2010), the mean fecundity in *Trichogramma* on *E. kuehniella* was at best around a hundred, and each of our host egg strips counted several hundreds of eggs, we can assume that our study was indeed done in a non-limiting context. Furthermore, in general, only one adult emerges from *E. kuehniella* eggs in the end (Corrigan et al., 1995; Klomp & Teerink, 1966).

Egg retention by refusing to oviposit was previously observed in *T. principium* and *T. brassicae* (Fleury & Boulétreau, 1993; Reznik et al., 2001, 1998). Therefore, egg retention may be present in all of the studied species and may affect fecundity measures in the timeframe of our experiment; see below for how this possibility was accounted for in the context of Data analyses.

In two-vial systems, effective dispersal (i.e. movement between patches leading to actual gene flow) was measured as a binary response, where one female is considered to have successfully dispersed if at least one parasitized egg was found on the strip present in the second plastic vial.

### Development time

After taking the pictures for fecundity, each isolated host egg strip was checked every day at around 9:00 a.m., 12:00 p.m., and 4:00 p.m. for the presence of emerged individuals. The development time of one replicate was considered to be the number of days between the female in the plastic vial starting to lay eggs and the emergence of the first offspring. Note that the true time is only known to a precision of two days, because of uncertainty in when precisely eggs were laid during the 48 h window after introduction in the system (see Data analyses for how this is accounted for).

### Data analyses

Data were then analysed with Bayesian multivariate multilevel/mixed models, using the brms R package, version 2.17.0 (Bürkner, 2017b), a front-end for the Stan language (Stan Development Team, 2022). The code was tested on R versions 4.1.2 to 4.2.0 (R Core Team, 2021).

The multivariate model architecture was made of four connected Generalized Linear Mixed sub-models, one for each response that are effective dispersal, development time, and fecundity divided into two components:

- Effective dispersal (the probability that the female successfully laid eggs in the arrival patch) was modelled with a Bernoulli distribution (with a logit link).
- Development time was modelled with a Log-Normal distribution, often chosen for time-to-event data. Because of the 48 h time period where the female was allowed to reproduce, development times were interval censored (with 48 h wide intervals);
- For fecundity, initial models showed evidence of both potential zero inflation and overdispersion, therefore a Zero-inflated Negative binomial distribution was used. This effectively separates the response variable into two components, “structural zeroes” and counts, each with a valid biological meaning (Blasco-Moreno et al., 2019):

– On one hand, the zero-inflated part of the distribution, similar to a Bernoulli model, modelled an excess of non-parasitized replicates compared to a negative binomial model. Given that egg retention is common in *Trichogramma* species, leading to delays in egg-laying of up to several days commonly (Reznik et al., 2001), a biologically plausible reading of these structural zeroes component is the probability of retention in the 48 h of the experiment;
– On the other hand, a Negative binomial component (with a log link) was interpreted as the fecundity of individuals that did not perform egg retention. From now on, we will use “fecundity without retention” to refer to this fecundity component (i.e. the effectively egg-laying individuals only), and “overall fecundity” will refer to the mean number of eggs laid by all individuals, including those potentially doing retention.

We used the model architecture described above for two multivariate models. The two multivariate models were fitted to observe how variance in traits and the covariance between traits are partitioned at the inter- and intra-specific levels. The first model incorporated both line and species-level effects, structuring the variance into intra- and inter-specific levels. The second model only had line effects as predictors, and therefore assumed that individuals from two conspecific lines do not resemble each other more than individuals from two randomly selected lines. In both cases, the same predictors were used for all four responses.

The first model included species-level effects as a fixed effect, mostly due to the low number of species studied, and line identity was coded as a random effect, while the second model only included line-level random effects. To account for line-level correlations between the response variables, line-level random effects for the two models were modelled as drawn from a shared variance-covariance matrix (Bürkner, 2017a).

While phylogenetic comparative methods could be used in this context, as some of the variations could be explained by shared ancestry (Felsenstein, 1985), there is no phylogenetic tree available for all lines used we could include (Hadfield & Nakagawa, 2010). Our first model, splitting variation into species and line components is nonetheless similar to the “taxonomic model” suggested in these cases where tree data are absent (Hadfield & Nakagawa, 2010).

The model formulas and the priors used (mostly weakly informative priors based on or modified from McElreath, 2020) are described in detail in **Supplementary Material S3.** The models were run with four chains during 4500 iterations each, with the first 2000 iterations for each chain used as warmup. This led to good chain convergence and sample size, as checked using the statistics proposed by Vehtari et al (2021). Model outputs were then checked using posterior predictive checks to compare predictions with empirical dataset (as suggested by Gabry et al., 2019). See the “Data and code availability” section for links to an archived version of the annotated model code.

## Results

*Trichogramma bourarachae* had lower fecundity without retention and higher development time than *Trichogramma brassicae*, while *T. semblidis* only had a lower development time than *T. bourarachae* but no clear difference in fecundity without retention (**Table 2**, **Figure 2B, C**). There were no other clear species differences (based on 95 % intervals of pairwise differences) in fecundity or development time. We did not find any evidence for between-species differences in effective dispersal probabilities (**Table 2**, **Figure 2A**). *T. brassicae* and *T. semblidis* both had higher egg retention probabilities than *T. bourarachae* (pairwise differences of 0.20 [0.00; 0.40] and 0.29 [0.05; 0.53] respectively, **Figure 2D**).

**Figure 2:**
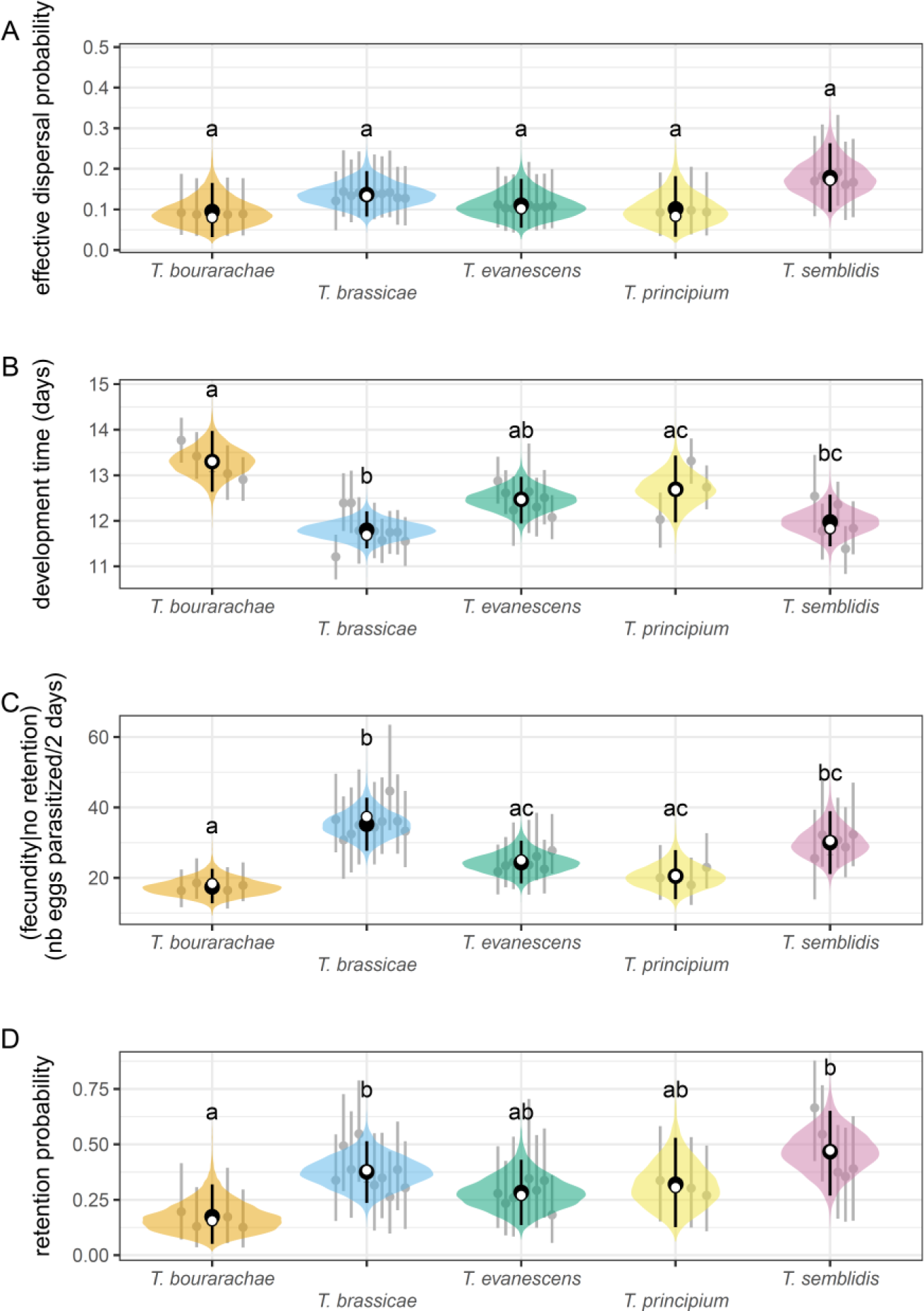
Posterior values as a function of the *Trichogramma* species for A) the probability of effective dispersal, B) the development time in days of the first offspring, C) the number of parasitized eggs for a single female when no retention occurred, and D) the probability for a female to perform egg retention during the experiment. 95 % posterior highest density intervals per line for each species are displayed in grey. Black dots represent posterior means and bars the 95 % intervals, while the posterior density distributions of fixed (species) effect predicted means are coloured depending on the species. For a given trait, two species with no index letters in common are considered to have “significant” pairwise comparison differences (i.e. the 95 % highest density interval of the difference does not include 0). White dots represent observed means per species, presented for illustrative purposes only (as they are calculated assuming all observed zeroes in egg numbers were attributable to retention, and using the midpoint of the 48 h interval for development time).

**Table 2:**
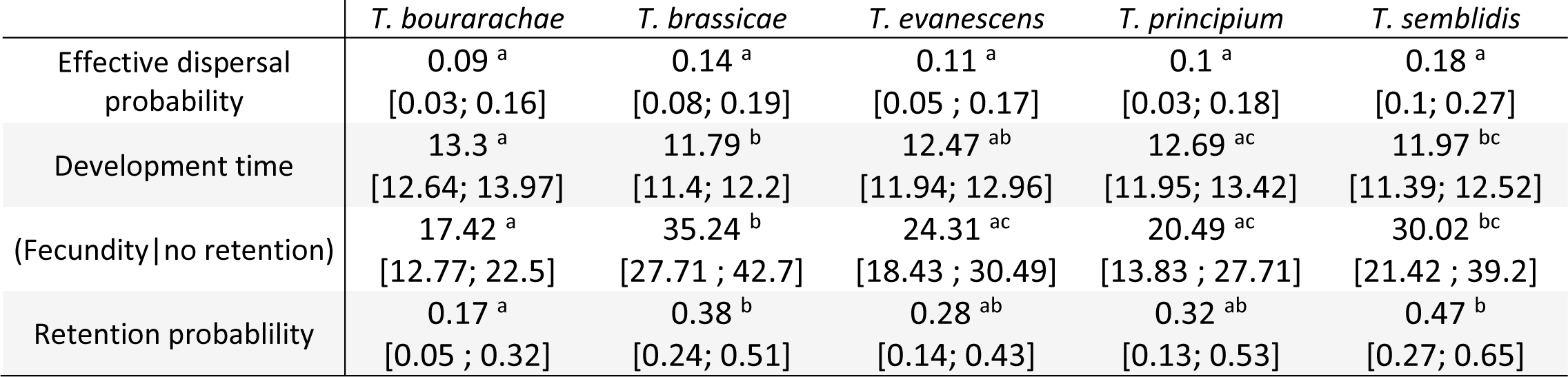
Mean posterior values and 95 % posterior highest density intervals per species, of single female fecundity in the absence of egg retention, said egg retention probability to occur during the experiment, the development time of the first offspring and effective dispersal probability. For a given trait, two species with no index letters in common are considered to have “significant” pairwise comparison differences.

|  | *T. bourarachae* | *T. brassicae* | *T. evanescens* | *T. principium* | *T. semblidis* |
| --- | --- | --- | --- | --- | --- |
| Effective dispersal probability | 0.09<sup>a</sup><br>[0.03; 0.16] | 0.14<sup>a</sup><br>[0.08; 0.19] | 0.11<sup>a</sup><br>[0.05 ; 0.17] | 0.1<sup>a</sup><br>[0.03; 0.18] | 0.18<sup>a</sup><br>[0.1; 0.27] |
| Development time | 13.3<sup>a</sup><br>[12.64; 13.97] | 11.79<sup>b</sup><br>[11.4; 12.2] | 12.47<sup>ab</sup><br>[11.94; 12.96] | 12.69<sup>ac</sup><br>[11.95; 13.42] | 11.97<sup>bc</sup><br>[11.39; 12.52] |
| (Fecundity\|no retention) | 17.42<sup>a</sup><br>[12.77; 22.5] | 35.24<sup>b</sup><br>[27.71 ; 42.7] | 24.31<sup>ac</sup><br>[18.43 ; 30.49] | 20.49<sup>ac</sup><br>[13.83 ; 27.71] | 30.02<sup>bc</sup><br>[21.42 ; 39.2] |
| Retention probability | 0.17<sup>a</sup><br>[0.05 ; 0.32] | 0.38<sup>b</sup><br>[0.24; 0.51] | 0.28<sup>ab</sup><br>[0.14; 0.43] | 0.32<sup>ab</sup><br>[0.13; 0.53] | 0.47<sup>b</sup><br>[0.27; 0.65] |

Correlations between traits at the line level were analysed through the random effect correlation/covariance matrix. In the first model, differences across species were modelled with a fixed effect, so they were not included in random effect correlations, while the second model included both species- and line-level random effects. Therefore, any qualitative difference between the two models can be interpreted as an effect at the species level.

The only detectable correlations among traits were between fecundity without retention and development time (**Table 3**, **Figure 3**). There was a negative correlation between these two traits at the line level in the model where species effects were not partitioned out (-0.62 [-0.92; -0.28], see **Table 3 bottom**, see also the overall pattern **Figure 3**). However, when looking at the model where species differences are partitioned out into fixed effects (**Table 3 top**), this random effect negative correlation mostly vanishes (-0.22 [-0.76; 0.38]). This reflects the fact that the overall correlation highlighted in **Table 3 top** is mostly driven by between-species differences in both fecundity and development time (see **Figure 2** and species averages in **Figure 3**).

**Figure 3:**
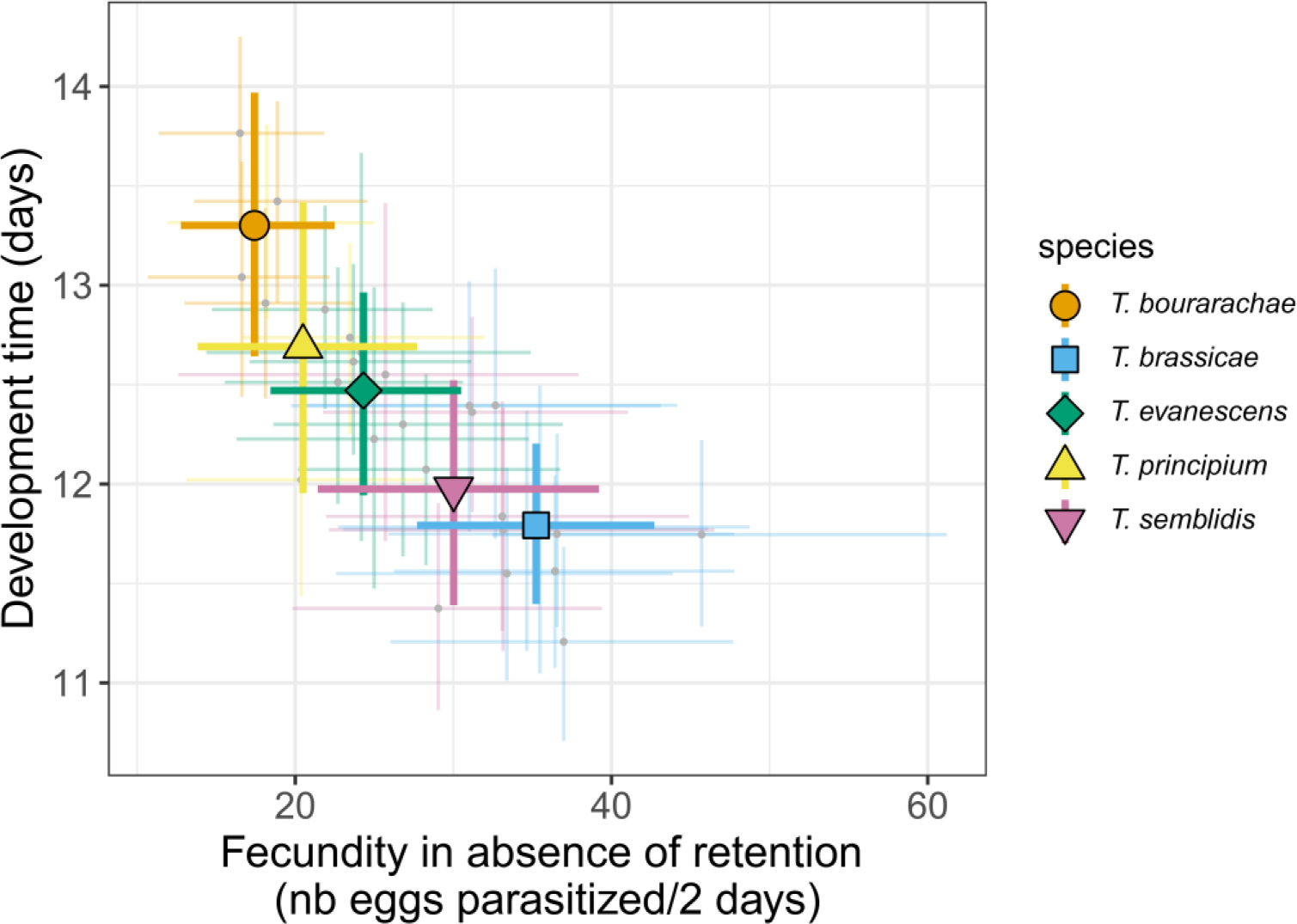
Posterior development time of *Trichogramma* species as a function of posterior fecundity in the absence of egg retention. Coloured crosses represent species 95 % posterior higher posterior density intervals for development time and fecundity, while coloured symbols represent species posterior means; line-level posterior means are displayed in grey and line-level posterior 95 % intervals are displayed in the colour of their corresponding species but more transparent.

**Table 3:** line-level random effect correlations among measured traits, represented by means and 95 % higher posterior density intervals. (top) Between-lines trait correlations, from the partitioned covariance model (species differences are excluded as fixed effects); (bottom) between-lines trait correlations from the model without fixed species effects. Intervals without 0 inside are presented in bold.

|  | effective dispersal probability | (Fecundity\|no retention) | Retention probability |
| --- | --- | --- | --- |
| **Within-species, among-line correlations (after excluding between-species differences as fixed effects)** |  |  |  |
| (Fecundity\|no retention) | 0.04 [-0.66; 0.72] |  |  |
| Retention probability | 0.04 [-0.65; 0.73] | -0.22 [-0.85; 0.4] |  |
| Development time | 0.12 [-0.58; 0.79] | -0.22 [-0.76; 0.38] | 0.2 [-0.37; 0.71] |
| **Overall among-line correlations, inclusive of between-species differences** |  |  |  |
| (Fecundity\|no retention) | 0.19 [-0.46; 0.82] |  |  |
| Retention probability | 0.15 [-0.52; 0.81] | 0.08 [-0.48; 0.61] |  |
| Development time | -0.08 [-0.71; 0.59] | **-0.62 [-0.92; -0.28]** | -0.17 [-0.64; 0.31] |

## Discussion

### Identification of one interspecific Pace-of-Life axis in Trichogramma

We found a negative between-line correlation between development time and fecundity in this subset of five *Trichogramma* species, with high fecundity without retention, fast development time on one side, and low fecundity, slow development on the other (**Figures 2, 3, Table3**). This correlation, which matches the classical pace-of-life axis (Healy et al., 2019) is mainly or only due to species-level differences: species with higher fecundity also had faster development times (**Figures 2, 3**), and the line-level correlation vanishes when species differences are partitioned out (**Table 3**). We note that even if there is no statistically significant correlation when the variance is structured within species and among lines, the sign of this correlation remains negative (**Table 3 top**), following a similar tendency to the interspecific negative correlation observed in **Table 3 bottom**. Having relatively similar patterns of interspecific and intraspecific correlation may result from close genetic correlations between development time and fecundity in *Trichogramma*, through pleiotropy or other strong genetic architecture, constraining the evolution of this trade-off among lines and species (Peiman & Robinson, 2017). It is also possible that at the metabolic level, resource acquisition and allocation may favour longer development times at the expense of fecundity or the opposite (Agrawal, 2020; Jørgensen & Fiksen, 2006; Stearns, 2000). The fact that a pace of life was found at the between-species level but not conclusively at the within-species level is in line with the existing literature, in which consistent pace-of-life axes are considered increasingly difficult to find as the taxonomic level narrows down, possibly due to scale-dependent mechanisms (Agrawal, 2020; Simons, 2002). Correlations at the species level act over a longer macroevolutionary timescale, where divergent positions on the pace-of-life axis of each species may represent long-term trade-offs and selection pressure over a wider environmental range than at lower levels, like lines. For lower levels, such as lines or individuals, traits may be more responsive to direct environment variation through phenotypic plasticity, and a shorter evolutionary timescale may lead to lower variation range compared to a higher level (Siefert et al., 2015).

However, this pace-of-life finding is based on splitting fecundity into what we interpret as egg retention and fecundity without retention components. While a significant negative correlation with development was found on the latter component of overall fecundity, results are more complex for retention probabilities. Indeed, there is no evidence for the line-level correlation between egg retention and other life-history traits (**Table 3**). Furthermore, at the species level, faster species (lower development time and higher fecundity in the absence of retention) were also the species with the highest retention probabilities (**Figure 2**, **Table 2**). If we interpret retention rates as a trade-off between present reproduction and future opportunities, then high retention can be seen as a “slow” trait; its association with “faster” life history traits may then appear paradoxical. It might be that fecundity in the absence of retention and retention probabilities are not actually separate traits, and that the trait correlations described above derive from their “artificial” separation by the statistical model. However, previous studies indicate that in *T. principium*, except for prolonged periods of egg-retention, individuals manifesting egg retention had similar fecundities in their first days of actual egg-laying and similar lifetime fecundities than individuals that did not (Reznik et al., 2001, 1998). Still in *T. principium*, there are indications that individuals manifesting egg retention have longer mean lifespans than individuals that immediately oviposit (Reznik et al., 2003; Reznik & Vaghina, 2007). These results support the idea that egg retention is a separate trait, interpretable as a mark of delayed reproduction (thus typically “slow” life history) rather than merely a component of reduced reproduction. While the pace of life remains an important and valuable structuring pattern in life histories, our results would agree with other studies showing that deviations from naïve expectations, where all traits should be either “slow” or “fast”, can be frequent (Bielby et al., 2007; Wright et al., 2020). In Wright et al (2020), their eco-evolutionary model presented possibly unexpected but existing life-history strategies, like “slow” adult reproduction alongside “fast” offspring survival (that the authors likened to an oak tree life history) or the opposite (represented by mayflies). However, because egg-laying was restricted to a 48 h window in our experiment, we cannot yet confirm this interpretation. Further studies measuring lifetime reproductive success, longevity or the way reproductive effort is spread throughout the lifetime may shed more light on the way life history is structured in *Trichogramma* wasps.

### No evidence for a syndrome linking effective dispersal probability and the pace of life

Effective dispersal probability varied the least among the four traits measured, with no evidence of between-species or even between-lines differences (**Figure 2**), and values were rather consistent with previous studies (Dahirel, Bertin, Calcagno, et al., 2021). There was also no correlation between effective dispersal and any of the other traits (**Table 3**), meaning that there is no evidence for dispersal/life-history syndromes at the line or species level in our set of *Trichogramma* species. Interestingly, this result on effective dispersal between patches completes previous studies on the activity of *Trichogramma* species. In Wajnberg & Colazza (1998), the authors showed a significant difference in the average area searched within one patch by *T. brassicae* isofemale lines while our results showed no differences in effective dispersal (**Figure 2**). In Reznik & Klyueva (2006), *T. principium* females manifesting egg retention had higher dispersal activity in a continuous environment than females that laid eggs beforehand. This discrepancy may be the result of a focus on different taxonomic levels: Reznik and Klyueva (2006)’s results deal with within-species and within-line covariation, versus between-lines and between-species in the present study. It may also result from differences in experimental designs and metrics used: the dispersal metrics used in Reznik and Klyueva (2006) are based on short-term (less than one day) and short-distance (up to 5 cm) movement on a continuous arena, compared to our experiment (two days and 40 cm between discrete patches). In that case, there may still exist in *Trichogramma* a pace-of-life syndrome linking life history to short-term activity and behaviour, but not effective dispersal. Indeed, correlations between short-term movement activity and life-history traits were also found in *T. evanescens* at the between-line level (Lartigue et al., 2022). While short-term activity metrics in uniform continuous environments are often considered valid proxies of longer-term dispersal between discontinuous patches (Pennekamp et al., 2019), this comparison of our study with the existing literature shows that this is not always the case. Dispersal is extremely context-dependent, including current resource availability (Fronhofer et al., 2018); there is furthermore evidence that correlations between dispersal and other traits can be altered depending on whether individuals disperse from high-resource or low-resource contexts (Cote et al., 2022), but also how density can have an impact on both dispersal behaviour and its evolution (Bitume et al., 2013; Clobert et al., 2009; Poethke et al., 2016; Poethke & Hovestadt, 2002).

### The potential implications of context-dependence and especially density-dependence

Building on this point, our study ignored this potential for context dependence, as every female tested for a given trait was tested under the same low-density conditions (alone in the experimental design with a non-limiting host supply). Dispersal syndromes (Bonte & Dahirel, 2017; Cote et al., 2022; Ronce, 2012) and also pace-of-life syndromes can be context-dependent. Behavioural types can be dependent of the dispersal status and predation risks (Bell & Sih, 2007; Cote, Clobert, et al., 2010). Pace-of-life syndromes and/or their constituent traits may also depend on resource acquisition through plastic responses (Laskowski et al., 2021; Montiglio et al., 2018) or quality. In *Trichogramma* for instance, host egg species and quality can influence life-history traits (Paul et al., 1981), and we used a substitution host in the present study. Recent works suggest that population density and density fluctuations, in particular, may also play a key role in shaping the presence of a pace of life in the first place (Wright et al., 2020) and its association with behaviours: fast individuals may have a higher reproductive rate in low-density contexts, but their lower intra-specific competition is a disadvantage when close to the carrying capacity of an environment, and therefore are more likely to disperse to escape to lower densities where this competition is lessened (Wright et al., 2019). This interaction between the pace of life and density may interact with the overall density dependence of dispersal (Clobert et al., 2009; Harman et al., 2020), altering syndromes linking dispersal and life history. Given that there is evidence for dispersal and/or fecundity being density-dependent in several *Trichogramma* species (*T. brassicae*; Dahirel, Bertin, Calcagno, et al., 2021; *Trichogramma achaeae*, *T. chilonis* and *T. euproctidis*, Zboralski et al., 2016), further studies including density dependence may lead to more generalizable insights about pace of life and dispersal in *Trichogramma*.

### Implications for biocontrol improvement and perspectives

While studies on trade-offs (Bennett et al., 2002; Reznik & Klyueva, 2006; Zboralski et al., 2016) or pace-of-life syndromes (Lartigue et al., 2022) already existed in small biocontrol agents including *Trichogramma*, our results provide new insights on between-species comparisons and the taxonomic scales at which trait variation is important. Some species like *T. evanescens*, *T. cacoeciae* (Sigsgaard et al., 2017) or *T. brassicae* (Özder & Kara, 2010) are already well used as biocontrol agents. In that context, a choice might be needed between maximizing one trait or a set of traits of interest at the expense of the others. For *Trichogramma*, while having fast-developing and high-fecundity individuals can be beneficial to quick intervention and a higher number of host eggs parasitized, they are reared and released mainly at high densities, (Consoli et al., 2010) densities for which individuals with longer development time might fare better against the intra-species competition (Wright et al., 2019). For inoculative releases, where a small population of biocontrol agents in the area of interest must establish itself and reach higher densities in further generations, both fecundity and competitive abilities are to be favoured for efficiency (Smith, 1996). Our results suggest that for some purposes, selecting different species might actually be more successful than attempting to select specific lines within one already used species.

## Data availability

Data and R code to reproduce all analyses presented in this manuscript are available on GitHub (https://github.com/CGuicharnaud/Trichogramma_POL_dispersal_2023) and archived in Zenodo (https://doi.org/10.5281/zenodo.7544901).

## Acknowledgements

We would like to thank the Biological Resources Center “Egg Parasitoids Collection” (doi.org/10.15454/AY4LMT) for providing the *Trichogramma* lines, training on how to rear them, and for contamination checks at the beginning and the end of the experiment. CG was supported by a PhD fellowship funded by the French Ministry of Higher Education, Research and Innovation (MESRI). The research leading to these results has received funding from the French Agence Nationale de la Recherche (project PushToiDeLa, ANR-18-CE32-0008). MD was funded under the European Union‘s Horizon 2020 research and innovation programme (Marie Skłodowska-Curie grant agreement 101022802) during this project.

## Author contributions

CG, GG, EV and MD designed the experiment. Lines rearing was done by LL and CG, while CG, GG, EB, LL and EV conducted the experiments in the lab. CG and LL collected data; CG and MD analysed the data. CG wrote the first draft; all authors were able to read, edit, and approve the manuscript.

