## Supplementary material for "Life-history traits, pace of life and dispersal among and within five species of *Trichogramma* wasps: a comparative analysis": Supplementary.html

- **Supplementary
  Material 1** - Detailed information for lines
  used
- **Supplementary
  Material 2** - Picture of the experimental
  design
- **Supplementary
  Material 3 –** Detailed description of statistical
  models
- References

- Chloé
  Guicharnaud, Géraldine Groussier, Erwan Beranger, Laurent Lamy, Elodie
  Vercken, Maxime Dahirel

### Supplementary Material for ”Life-history traits, pace of life and dispersal among and within five species of *Trichogramma* wasps: a comparative analysis”


#### **Supplementary Material 1** - Detailed information for lines used

**Supplementary Table S1-1**: Detailed information on
the lines used, with the country, department, GPS coordinates of
collection place and the date when samples were collected. Lines marked
with a \* were shared by the biocontrol firm BioLine as part of a
collaborative project (https://www.colbics.eu/Main-Results/Intraspecific-diversity-in-Trichogramma-brassicae),
but their precise geographic origin is kept confidential

| Species | Lines | Capture year | Country | longitude | latitude |
| --- | --- | --- | --- | --- | --- |
| *T. bourarachae* | ISA11967 | 2016 | France | 6.90 | 43.82 |
| *T. bourarachae* | ISA11969 | 2016 | France | 6.90 | 43.82 |
| *T. bourarachae* | ISA5544 | 2015 | France | 7.12 | 43.65 |
| *T. bourarachae* | ISA6646 | 2015 | France | 6.90 | 43.82 |
| *T. brassicae* | F5-8\* | 2013 | France | - | - |
| *T. brassicae* | F3-2\* | 2013 | France | - | - |
| *T. brassicae* | F3-9\* | 2013 | France | - | - |
| *T. brassicae* | F5-11\* | 2013 | France | - | - |
| *T. brassicae* | F5-12\* | 2013 | France | - | - |
| *T. brassicae* | F6-4\* | 2013 | France | - | - |
| *T. brassicae* | I2-16\* | 2013 | Europe | - | - |
| *T. brassicae* | I6-5\* | 2013 | Europe | - | - |
| *T. brassicae* | PR002 | 2015 | France | 5.43 | 43.59 |
| *T. evanescens* | BIO-XK | 2013 | France | 4.38 | 44.53 |
| *T. evanescens* | BIO-XE | 2013 | France | 6.88 | 43.59 |
| *T. evanescens* | BIO-XA | 2010 | France | 6.11 | 43.9 |
| *T. evanescens* | N-05 | 2016 | France | -0.78 | 43.49 |
| *T. evanescens* | Q-05 | 2015 | France | 3.58 | 45.67 |
| *T. evanescens* | HY-05 | crossing of three lines from Lartigue et al. (2022) made in 2019 | | | |
| *T. evanescens* | H-03 | 2016 | France | 4.93 | 44.98 |
| *T. principium* | 81a | 1975 | France | 6.13 | 43.12 |
| *T. principium* | ISA11235 | 2016 | France | 7.18 | 43.77 |
| *T. principium* | ISA11367 | 2016 | France | 7.19 | 43.74 |
| *T. semblidis* | FPV025 | 2015 | France | 4.88 | 45.95 |
| *T. semblidis* | PR007 | 2015 | France | 5.43 | 43.59 |
| *T. semblidis* | CVR065 | 2016 | France | 0.44 | 44.21 |
| *T. semblidis* | FPV034\_A | 2015 | France | 4.88 | 45.95 |
| *T. semblidis* | BL110 | 2016 | France | 0.89 | 45.2 |

#### **Supplementary Material 2** - Picture of the experimental design

**Supplementary Figure S2-1:** Picture of host eggs
strips used in a two-vial system. Parasitized host eggs are identified
by their blackened state compared to unparasitized host eggs.

#### **Supplementary Material 3 –** Detailed description of statistical models

The development times \(T\_{s,l,i}\),
overall fecundities \(F\_{s,l,j}\) and
effective dispersal probabilities \(D\_{s,l,k}\) for each observation \(i\), \(j\)
or \(k\) from each line \(l\) belonging to species \(s\) can be modeled using the following
multivariate model (note that due to interval censoring, \(T\_{s,l,i}\) is not observed directly, only
a 2-day wide interval \([t\_{s,l,i};t\_{s,l,i}+2]\) such that \(t\_{s,l,i}≤ T\_{s,l,i}≤t\_{s,l,i}+2\)): \[T\_{s,l,i} \sim \text{Log Normal}
(μ\_{s,l},σ\_{[T]}),\] \[F\_{s,l,j} \sim
\text{ZINegative-Binomial}(π\_{s,l},λ\_{s,l},φ),\] \[D\_{s,l,k} \sim
\text{Bernoulli}(p\_{s,l}),\]

Where \(π\) is the probability of
excess zeroes, which we interpret in this context as the retention
probability.

##### Model partitioning variance between species fixed effects and lines random effects

The models for each parameters are then as follows: \[μ\_{s,l}= β\_{[μ]s}+ α\_{[μ]l},\] \[\text{logit}(π\_{s,l})= β\_{[π]s}+
α\_{[π]l},\] \[\text{log}(λ\_{s,l})=
β\_{[λ]s}+ α\_{[λ]l},\] \[\text{logit}(p\_{s,l})= β\_{[p]s}+
α\_{[p]l},\]

The line-level random effects are distributed as follows: \[
\begin{bmatrix}
α\_{[μ]l}\\
α\_{[π]l} \\
α\_{[λ]l} \\
α\_{[p]l}
\end{bmatrix} \sim
\text{MVNormal}
\left(
\begin{bmatrix}
0\\
0 \\
0 \\
0
\end{bmatrix} ,Ω
\right)
\]

Where the line-level covariance matrix \(Ω\) can be decomposed into the random
effects standard deviations and the correlation matrix \(R\): \[
Ω = \begin{bmatrix}
\Large σ\_{α\_{[μ]}} & 0 & 0 & 0\\[0.3em]
0 & \Large σ\_{α\_{[π]}} & 0 & 0 \\[0.3em]
0 & 0 & \Large σ\_{α\_{[λ]}} & 0 \\[0.3em]
0 & 0 & 0 & \Large σ\_{α\_{[p]}}
\end{bmatrix}
R
\begin{bmatrix}
\Large σ\_{α\_{[μ]}} & 0 & 0 & 0\\[0.3em]
0 & \Large σ\_{α\_{[π]}} & 0 & 0 \\[0.3em]
0 & 0 & \Large σ\_{α\_{[λ]}} & 0 \\[0.3em]
0 & 0 & 0 & \Large σ\_{α\_{[p]}}
\end{bmatrix}
\]

We set weakly informative priors as suggested in McElreath (2020). We used a Normal (log(11), 1) and
Normal (log(19), 1) prior for fixed effect species intercepts \(β\_{[μ]}\) and \(β\_{[λ]}\) respectively. The prior means
here were shifted from the usual 0 to values based on the mean
development time and fecundity of *Trichogramma* in our dataset.
We used Normal(0, 1.5) priors for the fixed species effects \(β\_{[π]}\) and \(β\_{[p]}\), which are both interpretable as
the logit of a proportion. We used Half−Normal(0, 1) priors for all
standard deviations \(σ\) and a
LKJCorr(2) prior for the correlation matrix \(R\). Finally, for the negative-binomial
shape parameter \(φ\), we used a
Half−Normal (0, 1) on its inverse (1/\(φ\)), as recommended in https://github.com/stan-dev/stan/wiki/Prior-Choice-Recommendations.

##### Model with lines random effects only

The second model fitted is nearly identical to the one above, except
that there is no species-level fixed effects anymore, only overall
intercepts: \[μ\_{s,l}= β\_{0[μ]}+
α\_{[μ]l},\] \[\text{logit}(π\_{s,l})=
β\_{0[π]}+ α\_{[π]l},\] \[\text{log}(λ\_{s,l})= β\_{0[λ]}+ α\_{[λ]l},\]
\[\text{logit}(p\_{s,l})= β\_{0[p]}+
α\_{[p]l}.\]

The remainder of the model, including priors is otherwise
unchanged.

McElreath, Richard. 2020. *Statistical Rethinking: A Bayesian Course
with Examples in R and Stan*. 2nd ed. CRC Texts in Statistical
Science. Boca Raton: Taylor and Francis, CRC Press.
